# Salmonella injectisome penetration of macrophages triggers rapid translation of transcription factors and protection against cell death

**DOI:** 10.1101/2023.07.21.550113

**Authors:** George Wood, Rebecca Johnson, Jessica Powell, Owain J. Bryant, Filip Lastovka, Matt Brember, Pani Tourlomousis, John P. Carr, Clare E. Bryant, Betty Y.W. Chung

## Abstract

During bacterial infection both the host cell and its invader must divert intracellular resources to synthesise specific proteins in a timely manner. For the host, these factors may be needed for innate immune responses, including programmed cell death, and in the bacteria newly synthesized proteins may be survival factors needed to counteract host responses. *Salmonella* is an important food-borne bacterial pathogen that invades and multiplies within host cells. It is well established that invasion of epithelial cells is dependent upon the SPI-1 Type III injectisome, a biological needle that penetrates and secretes effectors into host cells to promote internalization. However, the importance of the SPI-1 injectisome in infection of professional phagocytes such as macrophages, which are the predominant host cell type during systemic infection, is less clear. Through time resolved parallel transcriptomic and translatomic studies of macrophage infection, we revealed that SPI-1 injectisome-dependent infection of macrophages triggers rapid translation of transcription factor mRNAs, including Early Growth Response 1 (*Egr1*). Despite the short half-life of EGR1 protein, its swift synthesis within the initial hour of infection is sufficient to inhibit transcription of pro-inflammatory genes and thereby restrain inflammatory responses and programmed cell death within the first hour of during early infection. This transient period of inflammatory suppression in macrophages is exploited by *Salmonella* to establish infection and sheds new insight on the importance of translational activation in host-pathogen dynamics during *Salmonella* infection.

## Introduction

The gene expression profiles of both host^1–5^ and pathogen^6^ are altered dramatically when they interact. This response is shaped by a co-evolutionary arms race where the host seeks to detect and counter the invading pathogen^3,5^, whilst the pathogen aims to evade this response^6^ and modify the host environment to better suit its survival and replication^1,2^. Changes in gene expression during infection are thus the net result of these competing goals, the balance of which can ultimately determine the infection outcome.

Many members of the Gram-negative bacterial genus *Salmonella* are facultative intracellular pathogens that infect a diverse spectrum of hosts. *Salmonella* species cause a range of diseases in humans, from typhoid fever to gastroenteritis. *Salmonella*’s ability to establish an intracellular infection is key to its pathogenesis^7^. During infection, the bacterium invades host cells, including the epithelial cells lining the intestinal tract and immune cells such as macrophages. Once internalized, *Salmonella* resides in a specialised membrane-bound compartment, the *Salmonella*-containing vacuole (SCV). The SCV provides a protective niche for the pathogen while giving it access to host cell nutrients to support its replication^8,9^.

The intracellular lifestyle of pathogenic *Salmonella* is supported by two Type III secretion systems (T3SS) with different substrate specificity. The SPI-1 T3SS, also called the SPI-1 injectisome (hereafter, ‘injectisome’) is a multi-protein complex that spans the bacterial inner and outer membranes and cell wall, and transports proteins into target cells along a homomeric needle-like structure that is inserted into the host cell membrane by the bacterial effectors SipB, SipC and SipD^10–12^. PrgJ forms the inner rod that connects the injectisome basal body embedded in the *Salmonella* envelope with the needle structure and thus acts as a channel bridging the bacterial and host cytoplasms^10^. In epithelial cells, the injectisome is important in aiding host cell invasion via the trigger mechanism^13–15^, though its role in establishing infection in professional phagocytes, such as macrophages, is less well understood. The SPI-2 T3SS is expressed once the bacterium is internalized and supports the intracellular lifecycle of the pathogen^8,16^.

To allow transport of effector proteins via the injectisome into the host cell, the *Salmonella* translocon subunits SipB and SipC are also secreted via the injectisome and inserted into the host cell plasma membrane to form a transient pore^11,12,17,18^. This insertion leads to transient loss of membrane integrity, therefore triggering an osmotic stress response and collapse of ion gradients with Ca^2+^ influx, and Cl^-^ and K^+^ efflux. Injectisome-dependent membrane damage is demonstrated by the haemolysin activity of the injectisome on red blood cells, which lyse upon SipBC insertion into their membranes^12^. While it is known that loss of membrane integrity, and the associated disruption of ion gradients, triggers inflammatory response pathways and cell death^17,19,20^, the importance of this damage in eliciting host responses to *Salmonella* remains unclear.

In systemic infection, macrophages are the predominant host cell type and *Salmonella* survival in macrophages has been reported to be critical for virulence^7,21^. This is somewhat paradoxical given the importance of macrophages for the detection and elimination of pathogens^22^. Indeed, macrophages poses many receptors that detect *Salmonella* pathogen associated molecular patterns (PAMPs), which are crucial in controlling infection. This includes toll-like receptors (TLRs) such as TLR4 which detects bacterial lipopolysaccharide (LPS) on the *Salmonella* cell surface^23^ as well as intracellular immune receptors that can activate the inflammasome and lead to inflammatory cell death^24–26^. In tissue culture, the majority macrophages die within the first few hour of infection^27–29^ and this rapid cytotoxicity is dependent on the injectisome^27,29^. The balance between macrophage survival and death will influence the outcome of infection^22^.

Although much research has been carried out on the transcriptional response of host cells to *Salmonella* infection^2–4^, the analysis of gene expression is incomplete without also exploring regulation at the level of translation, i.e. protein synthesis. This is of particular importance given the critical nature of events occurring very early in infection. Translational regulation has the potential to allow for rapid responses, either through modulating translation of pre-existing mRNAs and/or by enhancing translation of the newly transcribed mRNAs. Indeed, previous studies have identified potent translational upregulation of inflammatory genes, such as *Tnf*, in macrophages following stimulation of TLR4 with purified LPS^30,31^.

Here we utilized paired ribosome profiling and RNA-Seq over a time course of infection with wild type or SPI-1 injectisome mutant *Salmonella* in macrophages to understand the dynamics of gene expression regulation throughout injectisome-dependent infection. We identified that the translational response precedes the transcriptional response within sixty min post infection, and it is enriched for genes that encode DNA binding proteins. Within the translationally induced DNA binding proteins, we identified Early Growth Response 1 (*Egr1*). *Egr1* showed rapid injectisome-dependent transcriptional induction, with even greater translational induction, enabling the rapid and robust production of the EGR1 transcription factor early in infection. We further demonstrated that, while the EGR1 protein turnover is rapid, it acts as a longer term transcriptional suppressor of inflammatory genes triggered by *Salmonella* infection. We hypothesise that this creates an early window of opportunity for *Salmonella* to circumvent innate immunity, allowing it to successfully establish infection.

## Results

### *Salmonella* infection triggers rapid host cell responses

*Salmonella* entry into immortalized murine bone marrow derived macrophages (iBMDM, hereafter ‘macrophages’ unless otherwise stated) is markedly enhanced by the injectisome, although the mechanism in which it is involved in the invasion of professional phagocytes is unclear^13–15,32^. *Salmonella* cells associated with host cell membrane, i.e. actively invading bacteria, were seen as early as 5 min post-exposure, with *Salmonella* rapidly internalized within 15 min (Figure 1A-B). There was little further macrophage infection after 15 min of infection, indicating a potential change in susceptibility (Figure S1A). Injectisome-independent internalization was assayed through infection with mutant *Salmonella* unable to assemble the injectisome needle, generated by knockout of the SPI-1 inner-rod protein PrgJ^33^ and referred to as the ‘injectisome mutant’ hereafter. Invasion by injectisome mutant *Salmonella* was also observable within this timeframe, though to a lesser degree than for wildtype (WT) bacteria (Figure S1B).

**Fig 1:**
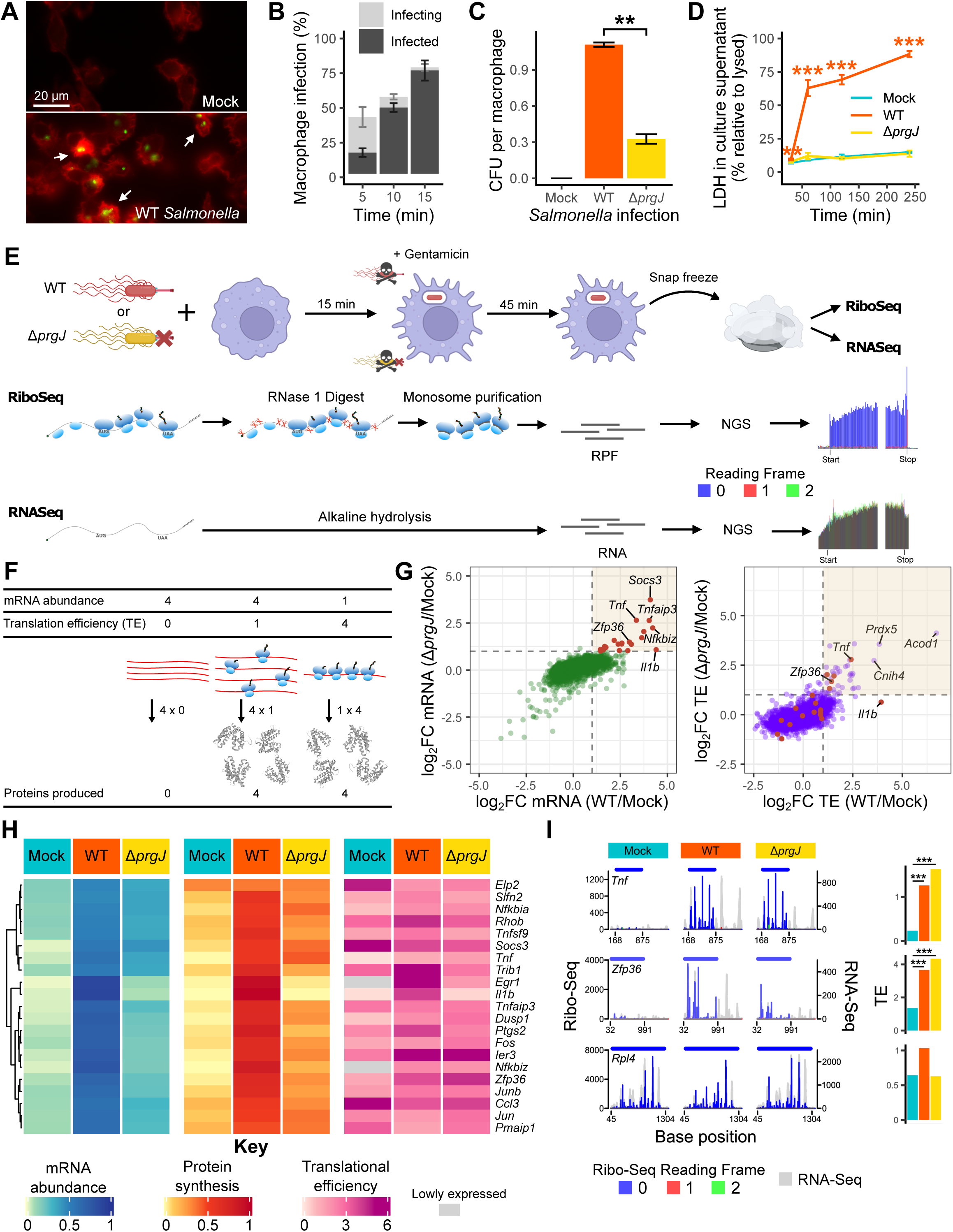
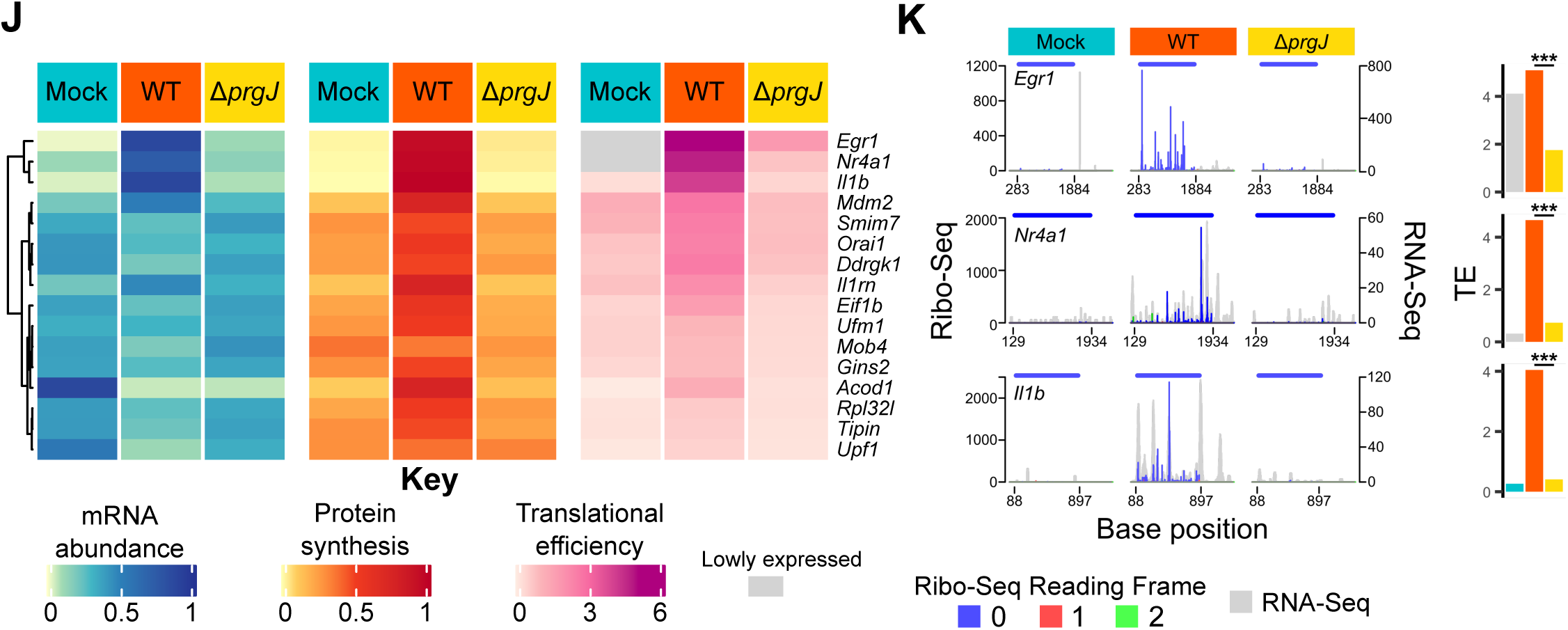
*Salmonella* SPI-1 dependent infection rapidly alters macrophage translation. (**A**) Mock infected and WT *Salmonella* infected macrophages 5 min post infection. Actin is shown in red, and *Salmonella* are shown in green. Arrows indicate *Salmonella* associated with the cell membrane. (**B**) Percentage of macrophages affected by WT *Salmonella* during the first 15 min of infection determined by immunofluorescent microscopy. Infected cells contain intracellular *Salmonella* infecting cells do not but have *Salmonella* associated with the host cell membrane, *n*=3. (**C**) *Salmonella* colony forming units (CFU) recovered 75 min post WT and SPI-1 deficient (Δ*prgJ*) *Salmonella* infection per host macrophage. Significance determined by Student’s t-test; *n*=2. (**D**) Cytotoxicity of WT or Δ*prgJ Salmonella* or mock infection of macrophages as determined by lactate dehydrogenase (LDH) release into culture supernatant. Significance determined by Students t-test; *n*=2. (**E**) Experimental outline detailing the preparation of ribosome profiling and RNA-Seq libraries from infected macrophages. Details are provided in the materials and methods (RPF: ribosome-protected RNA fragments, NGS: next-generation sequencing). (**F**) Diagram illustrating the relationship between mRNA abundance (measured by RNA-Seq), protein synthesis (measured by Ribo-Seq) and translational efficiency (TE). (**G**) Comparison of changes in mRNA abundance (left) and TE (right) on infection with WT or Δ*prgJ Salmonella* over mock infection. Genes upregulated transcriptionally in both (log_2_FC > 1) are shown in red. (**H**) Heatmap showing mRNA abundance, protein synthesis and TE of transcriptionally upregulated genes highlighted in G. Genes are ordered by hierarchical clustering of mRNA abundance across all conditions (left) (**I**) Ribo-Seq and RNA-Seq coverage of *Tnf* and *Zfp36* transcripts, which are known to have increased TE on exposure to bacterial PAMPs. Ribo-Seq reads are represented by their P site position and colored by their reading frame relative to the start codon of the coding sequence, represented by the bar above each plot with the start and stop positions indicated on the x-axis. *Rpl4* is presented as a control gene expressed in all samples. Significance determined using Xtail as described in the Materials and Methods (**J**) Normalized mRNA abundance, protein synthesis and TE of genes with log_2_FC in TE of WT over Δ*prgJ* infected macrophages greater than 1.5 and at least 50 normalized Ribo-Seq counts in WT infection. Genes are ordered by hierarchical clustering of TE across all conditions (left). Genes with low read counts (i.e. those with normalized RNA-Seq and Ribo-Seq counts less than 5) are considered lowly expressed and therefore TE cannot be reliably calculated. (**K**) Ribo-Seq and RNA-Seq transcript coverage of *Egr1*, *Il1b* and *Nr4a1* as in **G**.

As previously described^27–29^, infection of macrophages with WT *Salmonella* led to rapid cell death within the first 60 min of infection. However, approximately 25% of macrophages survived despite the presence of viable intracellular bacteria (Figure 1C-D, S1A). In contrast, and similar to previous studies^34^, infection with injectisome mutant *Salmonella* did not induce any increase in macrophage cell death (Figure 1D).

### Injectisome-dependent infection leads to both transcriptional and translational induction of Early Growth Response 1 (EGR1) accumulation

As a significant proportion of macrophages survive the lethal effect of injectisome penetration (Figure 1A-D), we hypothesized that survival of infected cells beyond 60 min post-infection with WT *Salmonella* may be a consequence of rapid gene expression responses occurring within the first hour. This may be mediated by *de novo* transcription of mRNAs, which has been the focus for most previous studies^2–4^. However, we suspected the involvement of translational responses, which remain much less well understood, where the rate of protein synthesis (translational efficiency) from mRNAs may be upregulated to increase protein abundance more rapidly than could occur *via* transcription alone. Such an acceleration of protein synthesis could be of critical importance given the short survival timeframe of most infected cells. To investigate rapid transcriptional and translational changes in gene expression, parallel global transcriptomic (RNA-Seq) and translatomic (i.e., ribosome profiling, Ribo-Seq hereafter) analyses were performed on macrophages at 60 min following infection with either WT *Salmonella* or the injectisome mutant (Figure 1E, S1C).

Ribo-Seq is a highly sensitive method that reveals the global translatome at the time of harvest^35^. The technique determines the position of ribosomes by exploiting the protection from nuclease digestion of a discrete fragment of mRNA (∼30 nucleotides) conferred by elongating ribosomes. Deep sequencing of these ribosome-protected fragments (RPFs) generates a high-resolution view of the location and abundance of translating ribosomes on different mRNA species, reflecting the amount of synthesis of specific proteins (Figure 1E). In addition, while RNA-Seq enables quantification of total mRNA abundance, parallel RNA-Seq and Ribo-Seq enables quantification of translation efficiency, a measurement of how well each mRNA is being translated as distinct from total protein synthesized (Figure 1F).

The resolution of our Ribo-Seq data is high, as evident from the metagene analysis constructed with the software program riboSeqR^36^. The metagene translatome is a summed plot of all translated mRNAs, with weighted average number of nucleotide reads around the annotated coding start and stop sites, confirming accurate capture of the elongating ribosome movement as almost all Ribo-Seq reads overwhelming maps to the first codon position (S1C). This is further evidenced by our ability to directly visualize translation of single genes at remarkable accuracy (Figure 1I), as well as non-canonical translation events. For example, translation of *Atf4,* which is modulated by translation of two small upstream open reading frames (uORFs) embedded within the 5’UTR of ATF4^37^ (Figure S1E). The high-resolution nature of our data therefore enables accurate quantification of protein synthesis (i.e. total Ribo-Seq) and translational efficiency when combined with parallel RNA-Seq (Figure 1F).

Initial analysis of macrophages infected by both WT and injectisome mutant *Salmonella* identified many genes known to be upregulated upon exposure to bacterial PAMPs^30,31,38^, such as *Tnf* and *Zfp36*. These genes were upregulated not only in transcript abundance but also at the translational level (Figure 1G, H, I and S1D and F). For both of these genes, transcription was induced in response to either the WT or injectisome mutant *Salmonella*. Potent translational upregulation was also seen, i.e. enhanced translational efficiency resulting in greater protein synthesis than can be explained by an increase in *de novo* mRNA synthesis alone (see Figure 1F), as has been previously reported in macrophages exposed to purified bacterial LPS^30,31^.

Following this, transcripts that were subject to specific injectisome-dependent translational upregulation were investigated (Figure S1F). The key inflammasome component *Nlrp3* was among the selectively induced genes (Figure S1G). The *Nlrp3* transcript has recently been described to encode a uORF^39^ which can be readily visualised in this data (Figure S1H). The NLRP3 is activated in *Salmonella* infection but the uORF-mediated translational regulation of *Nlrp3* is currently unclear^33,40,41^. To further identify injectisome-dependent translationally regulated genes, transcripts where the log_2_ fold-change (log_2_FC) in translational efficiency was greater than 1.5 in WT *Salmonella* infected macrophages over macrophages infected by the injectisome mutant were selected (Figure 1J). Amongst the most translationally upregulated mRNAs were those encoding Early Growth Response 1 (EGR1), NR4A1 and Pro-Interleukin-1 beta (pro-IL-1β) (Figure 1J-K).

Interleukin-1 beta (IL-1β) is processed from its precursor pro-IL-1β into its mature, active form by caspases; proteases that are themselves activated by the inflammasome^26,42^. This precursor, encoded by *Il1b*, had a much greater translation efficiency in macrophages infected by WT *Salmonella* than the injectisome mutant (Figure 1J-K). The injectisome is known to transport bacterial effectors into the host cytosol that activate the inflammasome, including components of the translocon such as SipB/C^43,44^, consistent with our cytotoxicity assay (Figure 1D), leading to IL-1β production^25,45^. These data suggest that IL-1β precursor production is supported not only by increased mRNA transcription, but also post-transcriptionally by specific upregulation of translation of its mRNA. Previous reports describe the need for two signals to produce IL-1β: one to induce transcription of the precursor and another to activate the inflammasome^43,46,47^. Importantly, however, our data additionally reveal a strong role for translational upregulation in precursor production to rapidly facilitate overall IL-1β precursor protein production. Therefore, the injectisome makes an important contribution to macrophage pro-IL-1β production within the first 60 min, in addition to delivering the factors that stimulate pro-IL-1β cleavage leading to IL-1β production.

Both *Egr1* and *Nr4a1* are known immediate early genes that are rapidly transcribed in many cell types within minutes in response to a range of cellular stresses^48–50^. Here we show that while transcripts for both genes are almost absent in uninfected macrophages, rapid expression of these genes is enhanced by specific transcriptional upregulation leading to a surge of translational activation following injectisome-dependent infection (Figure 1J-K).

NR4A1, also known as NUR77, is involved in macrophage responses to proinflammatory stimuli. NR4A1 limits inflammation in models of sepsis and colitis, likely through antagonism of the NF-κB pathway^51,52^. More recently, however, it has been reported to increase expression of proinflammatory cytokines in mice infected with *Klebsiella pneumoniae*^53^. The role of NR4A1 is therefore certainly immunomodulatory but likely differs by cell type and context.

Of the three transcripts chosen for detailed study, the most translationally induced is the mRNA for EGR1 (Figure 1J-K). EGR1 is a zinc-finger family transcription factor that binds GC-rich consensus sequences in gene promoters and enhancers, and it can either activate or suppress transcription. Targets of EGR1 span diverse biological processes including immune responses, cell growth and differentiation, and cell death^54–59^. Due to its highly injectisome-dependent translational upregulation, the biological function of EGR1 in *Salmonella-*macrophage infection was further characterized in this study.

### Specific upregulation of EGR1 protein accumulation during the macrophage-*Salmonella* interaction is highly injectisome-dependent

Transcriptional upregulation of *Egr1* has previously been shown to be largely dependent on bacterial secretion systems in other infection contexts^2,60–62^. However, EGR1 has not been studied in the context of *Salmonella* infection of macrophages nor has post-transcriptional regulation of *Egr1* gene expression been studied. We were particularly interested in EGR1 given its association with cell death^54^ and its importance in macrophage development^49,57^.

At 60 min post-infection there was a clear increase in both transcription and translation of *Egr1* in WT-infected cells (Figure 1K, l and 2A). The increase in *Egr1* translation cannot be explained by the greater transcript abundance alone, but rather there is also an increase in *Egr1* mRNA translation efficiency. The translational efficiency of *Egr1* mRNA is three times higher in WT-infected cells compared to cells infected with the injectisome mutant (Figure 1K, l and 2A). In contrast, the *Egr1* expression in cells infected with the injectisome mutant were not much greater at 60 min post infection than in mock-infected cells. Reverse transcription-coupled quantitative PCR (RT-qPCR) for *Egr1* mRNA and immunoblot for EGR1 protein accumulation showed that during an infection time-course the upregulation was rapid but transient and that the half-lives of both *Egr1* mRNA and its protein product are short. While *Egr1* mRNA abundance peaked at 60 min and returned to baseline levels by 120 min (Figure 2B and C), the increase in protein abundance measured by immunoblotting was, as expected, offset from the increase in mRNA, peaking at 120 min post-infection and returning to undetectable levels by 240 min. The absence of EGR1 protein 240 min post-infection indicates a short half-life for EGR1 protein and that its biological effect is likely rapid (Figure 2C-D). In contrast, infection with the injectisome mutant led to a slight increase in *Egr1* mRNA abundance at 60 min, followed by a detectable increase in EGR1 protein by 120 min, with both mRNA and protein at considerably lower levels than in WT *Salmonella* infected cells (Figure 2A-D).

**Fig 2:**
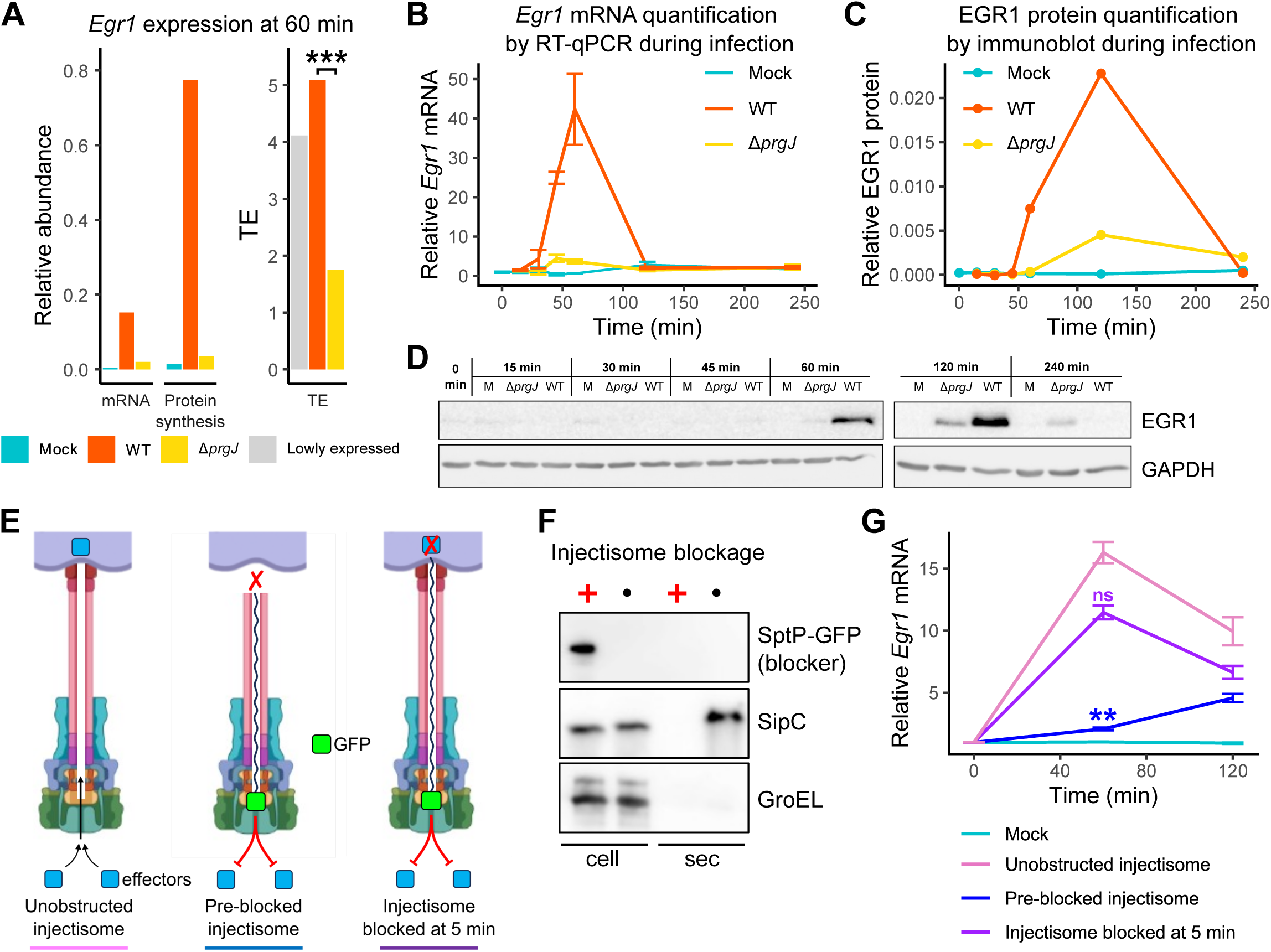
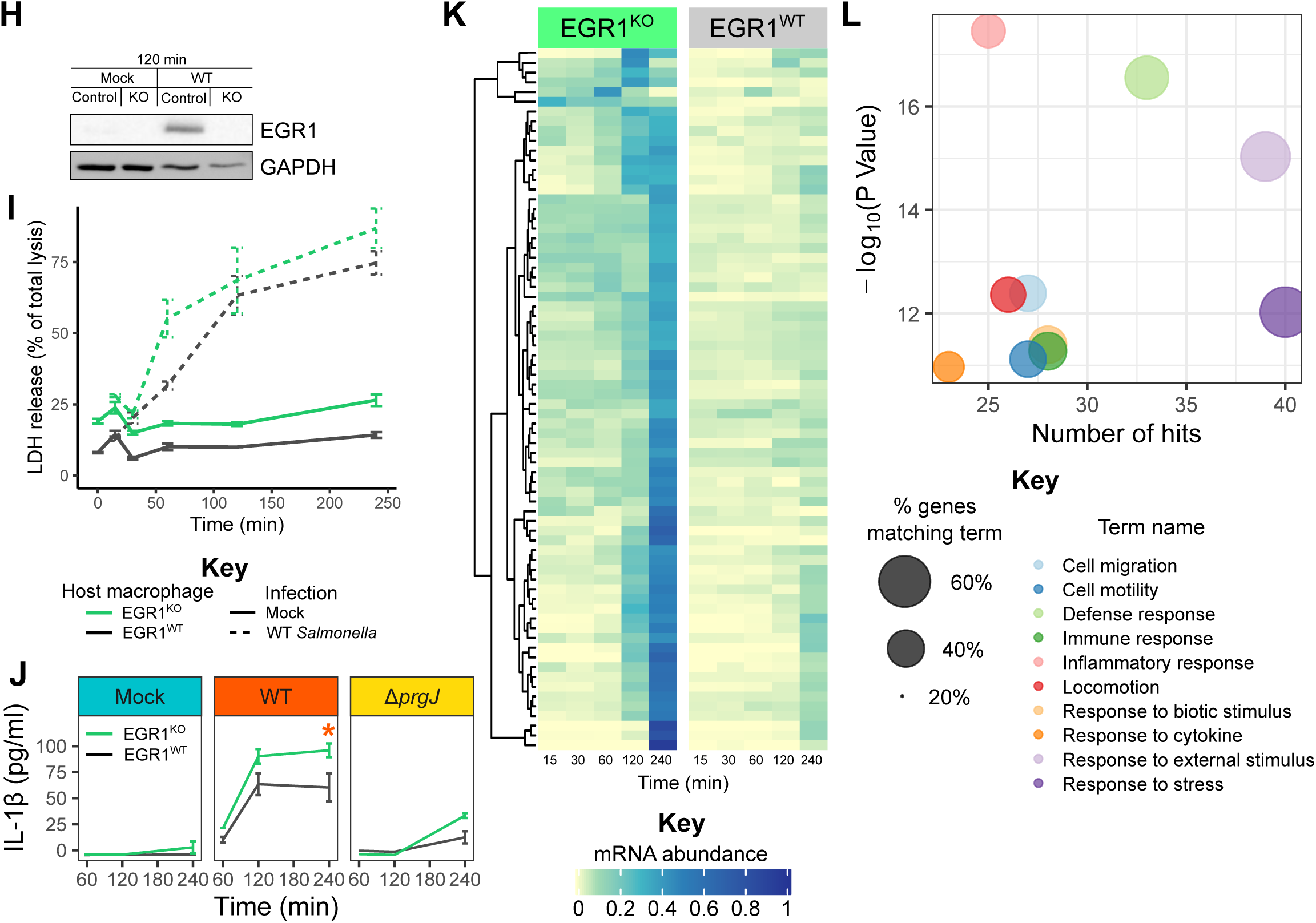
*Egr1* is rapidly induced in *Salmonella* infection to restrict expression of immune response genes. (**A**) mRNA abundance, protein synthesis and translational efficiency (TE) of *Egr1* from Fig 1K. For conditions with low *Egr1* expression TE cannot be reliably calculated (grey bar); significance determined using Xtail as described in the Materials and Methods. (**B**) *Egr1* transcript abundance over WT and SPI-1 deficient mutant (Δ*prgJ*) infection, normalized to the housekeeping gene *Supt16* and relative to cells prior to infection, *n*=2. (**C**) Quantification of EGR1 abundance from **D** normalized to GAPDH. (**D**) Immunoblot following EGR1 and GAPDH protein abundance across WT and Δ*prgJ* infection. An equal amount of cellular protein was loaded per lane. (**E**) Experimental outline illustrating the effect of the SptP-GFP injectisome blocking substrate on protein export via the injectisome. *Salmonella* cells not producing the blocking substrate can transport effector proteins into host cells (unobstructed injectisome, left). *Salmonella* cells expressing the injectisome blocking substrate before infection are unable to transport effector subunits via the injectisome (pre-blocked injectisome, middle). *Salmonella* cells were also incubated with macrophages for 5 min before inducing expression of the injectisome blocking substrate such that injectisomes can engage with macrophage cells but secretion of effectors proteins from *Salmonella* into macrophages are blocked at 5 min (injectisome blocked at 5 min, right). (**F**) Secretion analysis of WT *Salmonella* either expressing the SptP-GFP injectisome blocking subunit or carrying empty vector. Whole cell (cell) and secreted proteins (sec) from late-exponential-phase cultures were separated by SDS-PAGE and immunoblotted with anti-Myc-tag (SptP-GFP), anti-SipC or anti-GroEL antisera. (**G**) *Egr1* transcript abundance over the *Salmonella* infection time course with blockade of the SPI-1 injectisome induced at various timepoints. Significance determined by one-sided Student’s t-test; *n*=2. (**H**) Immunoblot showing EGR1 expression in EGR1^WT^ and EGR1^KO^ macrophages infected with WT *Salmonella* at 120 min post infection. (**I**) Cytotoxicity of WT or mock infection of EGR1^WT^ and EGR1^KO^ macrophages as determined by lactate dehydrogenase (LDH) release into culture supernatant. (**J**) IL-1β concentration in culture supernatant from infected EGR1^KO^ and EGR1^WT^ macrophages. *n*=2 except for 240 min WT *Salmonella* infection where n=3 for EGR1^WT^ and 4 for EGR1^KO^; significance determined using one-sided Student’s t-test. (**K**) Normalized mRNA abundance of transcripts upregulated (log_2_FC > 2) in EGR1^KO^ when compared to EGR1^WT^ macrophages at any timepoint in WT *Salmonella* infection. Genes are ordered by hierarchical clustering (left). (**L**) The 10 most significantly enriched gene ontology biological process terms in genes identified in J.

We reasoned that both *Egr1* transcriptional stimulation and the potent translational induction (i.e. higher translational efficiency) could be either a result of the direct interaction of the injectisome with macrophage, through insertion of the SipB/C translocon complex into the macrophage plasmalemma^18^, or due to transport of effector proteins once the injectisome is fully assembled and is in a secretion competent state. To distinguish between these possibilities, we engineered an injectisome blocking substrate in which the effector protein SptP is C-terminally fused to the green fluorescent protein (SptP-GFP). This injectisome blocking substrate is targeted to the export machinery and stalls within the injectisome export channel because the folding of the GFP moiety (unlike the effector protein sequence) is irreversible. Stalling occurs after needle assembly is complete, thereby blocking transport of effectors and translocon subunits through the injectisomes that have assembled on the bacterial cell surface^63^, preventing penetration of macrophage plasmalemma (Figure 2E-F). Obstructing the injectisome enabled us to uncouple the effect on *Egr1* upregulation upon penetration of macrophage plasmalemma by the injectisome from transport of effector and translocon subunits.

Expression of the SptP-GFP injectisome blocking substrate was controlled in the following manner: (1) uninduced, therefore recapitulates WT infection where the injectisome penetrates macrophages and delivers effectors; (2) Inducing expression of the blocking substrate 2 h prior to macrophage infection, resulting in assembly of injectisomes that are blocked with the SptP-GFP blocking substrate, preventing delivery of effectors whilst at the same time preventing delivery and insertion of translocon subunits (SipB/SipC) into the host cell membrane, which consequently abolishes penetration of macrophage plasmalemma; and (3) blockage of injectisome induced at 5 min post-infection, therefore allowing injectisomes to penetrate macrophage plasmalemma while inhibiting further delivery of effectors (Figure 2E). As expected, transcription of *Egr1* was upregulated when macrophages were infected by *Salmonella* with unobstructed injectisome (uninduced), but expression is significantly impaired in macrophages challenged by *Salmonella* with a pre-blocked, translocon-defective injectisome. However, *Egr1* transcript accumulation was only slightly reduced when penetration of injectisome is established but delivery of effector is prevented by SptP-GFP induction at 5 min post-infection (Figure 2G). Taken together, these results suggest that the trigger for overall *Egr1* protein accumulation occurs very rapidly during infection and is likely a direct result of injectisome penetration with the macrophage plasmalemma or the action of the first few effector molecules that make it through the injectisome.

### EGR1 protein restrains macrophage inflammatory responses to *Salmonella* infection

To further investigate the role of EGR1 during injectisome dependent infection of macrophages, we generated EGR1 knockout (EGR1^KO^) macrophages using CRISPR-Cas9 and confirmed the absence of EGR1 protein 120 min after infection with WT *Salmonella* (Figure 2H). Mutation of the *Egr1* coding sequence resulting in EGR1 knock-out in EGR1^KO^ macrophages was confirmed through genomic DNA sequencing (Figure S2A). Loss of the ability to produce EGR1 resulted in greater cell death at baseline, and this was further increased when infected by WT *Salmonella*, particularly between 30 and 120 min post infection (Figure 2I). This suggests EGR1 may play a role in limiting injectisome-induced macrophage death. Following this, the role of EGR1 in the inflammatory response was assessed by measuring the level of IL-1β produced by EGR1^KO^ and EGR1^WT^ macrophages during infection. This revealed significantly greater upregulation in EGR1^KO^ macrophages infected with WT *Salmonella*, confirming that EGR1 has a suppressive role in the inflammatory response.

As EGR1 is annotated as a DNA-binding protein, we hypothesized that EGR1 suppresses inflammation through transcriptional regulation and so time-resolved transcriptomic analysis (RNA-Seq) of WT *Salmonella-*infected EGR1^KO^ or EGR1^WT^ macrophages from 15 to 240 min post infection was performed. Principal component analysis of gene transcript levels showed a clear separation of samples by cell line and time post-infection. Principal component (PC) 1 largely reflects variation between the EGR1^KO^ and EGR1^WT^ cell lines, whereas PC2 separates the 120 min and 240 min infected samples from the 15 to 60 min infected samples and 15 to 240 min mock inoculation treatments (Figure S2B). We observed a significant increase in the abundance of transcripts associated with immune responses in the EGR1^KO^ macrophages compared to the EGR1^WT^ macrophages, particularly at 240 min post-infection, confirming the suppression of transcription of inflammatory genes by EGR1 during *Salmonella* infection (Figure 2K-L; Table S1-2). This includes *Il1b*, which showed greater upregulation of transcription in EGR1^KO^ macrophages infected with WT *Salmonella*. This is likely the cause of the greater IL-1β secretion during *Salmonella* infection in the absence of EGR1 (Figure 2J) and confirmed EGR1 as an important transcriptional immunomodulator. In addition, gene ontology enrichment analysis also revealed significant upregulation of known pro-cell death genes in infection of the EGR1^KO^ macrophages, including the FAS death receptor, consistent with a role of EGR1 in limiting macrophage death (Figure 2L; Table S1-2).

### The transcriptional and translational dynamics of Salmonella during macrophage infection

We have demonstrated that injectisome-dependent infection triggers surges both in *de novo* mRNA synthesis and in translational efficiency of specific host mRNAs such as *Egr1*, leading to rapid but transient accumulation of EGR1 protein. To assess global expression dynamics throughout *Salmonella* infection, we therefore performed time-resolved parallel RNA-Seq and Ribo-Seq of primary bone marrow-derived macrophages infected with WT *Salmonella* or the injectisome mutant (Figure 3A). The use of primary macrophages (rather than the iBMDMs used in previous experiments) allows us to compare injectisome-*versus* PAMP-dependent responses in a system that more closely resembles *in vivo* infection. The primary macrophages showed similar rates of infection to iBMDMs for WT *Salmonella*, but a greater proportion were infected with the injectisome mutant compared to iBMDMs (Figure S1B and S3A).

**Fig 3:**
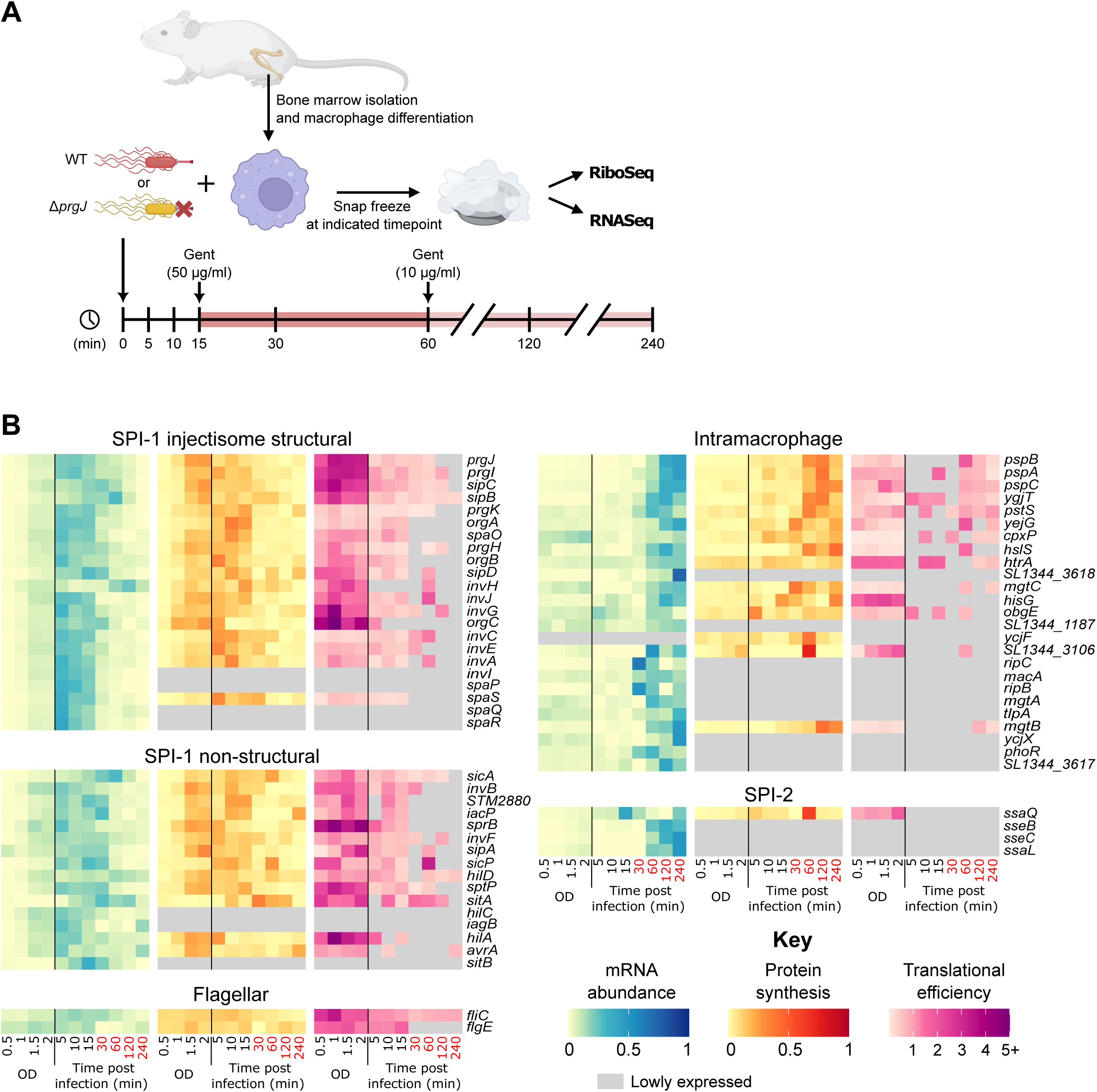
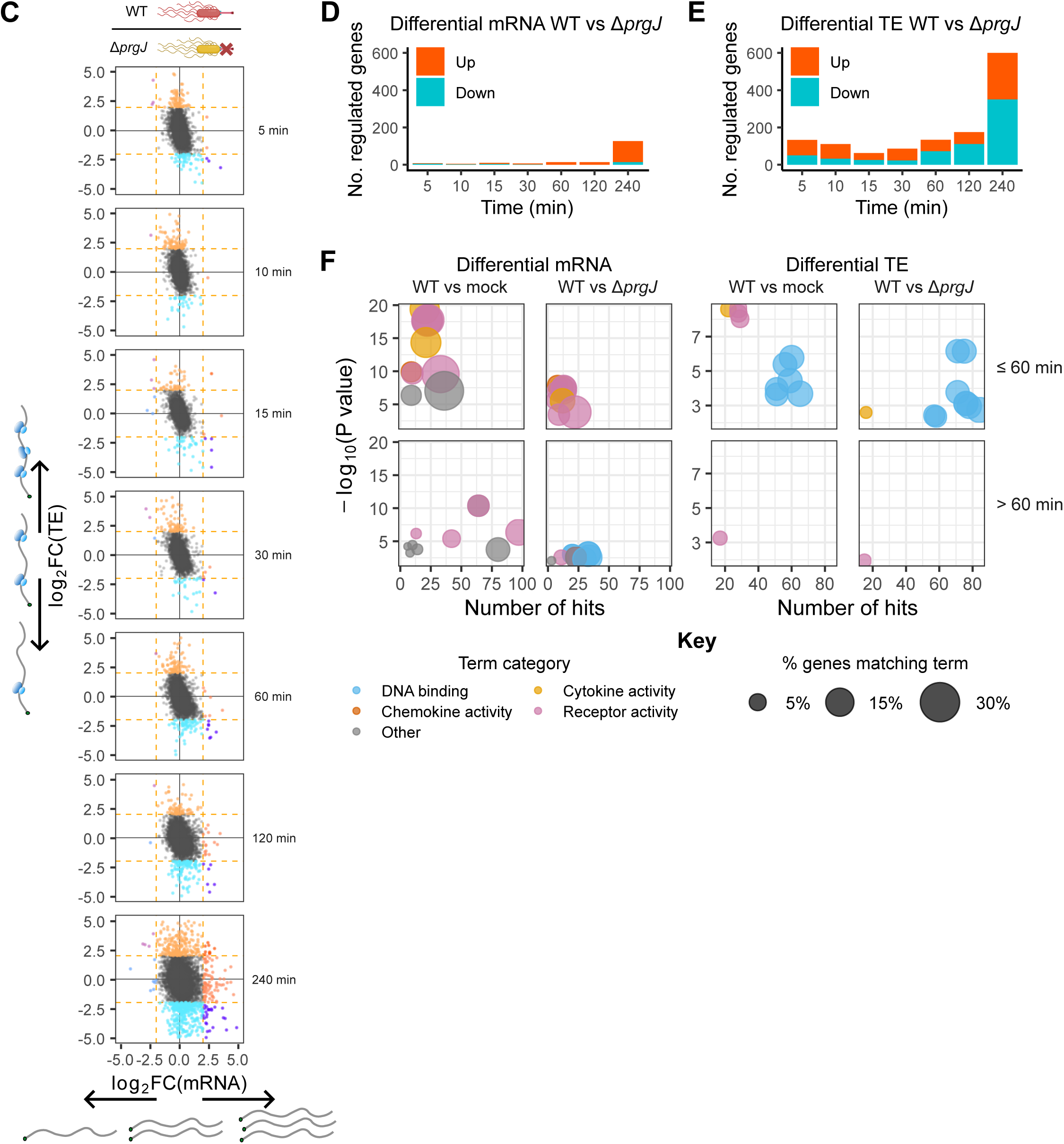
Time resolved RiboSeq reveals dynamic changes in host and bacterial translation. Data from two replicates. (**A**) Outline of infection time course. Primary bone marrow-derived macrophages were generated from mice and infected with WT or Δ*prgJ* mutant *Salmonella*. Gentamicin was used to kill extracellular bacteria at 15 min and its concentration was reduced after 1 h. (**B**) Transient expression dynamics of select WT *Salmonella* genes at different optical densities (OD) and over 4 h of macrophage infection, at the level of mRNA abundance, protein synthesis and translation efficiency. (**C**) Differential regulation of host gene expression as measured by the log_2_FC of translational efficiency (TE) vs log_2_FC mRNA abundance in WT *Salmonella* infection over Δ*prgJ* infection. Dashed orange lines show the log_2_FC cutoffs (±2) used to select differentially expressed genes, and genes that pass these thresholds are colored. Only genes with log_2_FC in mRNA and TE between -5 and 5 are shown here (see Fig S4B for uncropped plots). (**D** and **E**) Number of genes differentially expressed between WT and Δ*prgJ Salmonella* infection at both the TE (**D**) and mRNA (**E**) levels. (**F**) Top 10 enriched GO molecular function terms in differentially expressed genes at both the TE (right) and mRNA (left) levels. Genes were split by when they were differentially expressed: at or before 60 min, and after 60 min post infection.

Due to the nature of the mRNA enrichment and library preparation method employed, both host and bacterial translational and transcriptional dynamics were simultaneously captured over the course of infection. Transcription and translation were also assessed in *Salmonella* grown in Luria Broth (or lysogeny broth, LB) at various optical densities (OD). It has been established that expression of SPI-1 in LB culture peaks at late exponential phase, OD 2^29^, as they were prepared for in these infection assays. This data demonstrates this increase in expression of SPI-1 injectisome structural genes as the bacteria approach OD 2, through enhanced translation (Figure 3B). Remarkably, we see significant transcriptional induction of SPI-1 genes upon contact with macrophages, particularly structural components such as Prg I/J and K, as early as 5-15 min. aligning with the timeframe of the initial Salmonella-macrophage interaction (Figure 1A-C). This indicates that *Salmonella* respond to the proximity of macrophages by upregulating expression of certain genes that enable invasion (Figure 3B), and suggests that rapid assembly of injectisomes occurs on the bacterial cell surface during the first 15 min. Indeed, the rapid assembly of additional T3SSs upon contact with host cells has also been suggested in *Yersinia enterocolitica* infection^64^. The transcriptional and translational response of non-structural SPI-1 genes that encode effector proteins, structural genes with a dual-role as effectors such as *SipB* and *SipC*, and other virulence factors also increased during the first 15 min whereas some genes, such as for *AvrA* and *IacP*, were maximally expressed between 2-4 h post infection (Figure 3B).

Notably, during the early stages of infection (within the first 30 min), we also observed increases in the translation efficiency of a subset of intramacrophage genes, including *htrA*, *hisG* and *cpxP* (Figure 3B). Intramacrophage genes play a crucial role in the survival and replication of the bacterium inside macrophages and in general maximal transcription for these genes occurs later. The functions of these genes include roles in magnesium and phosphate transport and the envelope stress response^65,66^. At the 30 min and 60 min time points, when *Salmonella* is intracellular and after SCV acidification^67^, the transcript abundance of SPI-2 genes increases while the abundance of the majority of SPI-1 transcripts decreases (Figure 3B). This agrees with the recognized switch of *Salmonella* secretion mediated by the transcriptional regulator SsrB. Concurrently we observed decreases in expression of the flagella components, consistent with previously reported SsrB-mediated repression^65,68,69^ (Figure 3B).

### A global translational response precedes transcriptional responses in macrophages following injectisome penetration

Analysis of parallel Ribo-Seq and RNA-Seq throughout the infection time course provided a global perspective of the dynamics of host gene expression at multiple levels, notably changes in RNA abundance and translation efficiency, by comparing primary macrophages infected with either WT *Salmonella* or the injectisome mutant with mock infection (Figure S4A). Within 5 min post infection, we can readily visualize that both WT and injectisome mutant *Salmonella* induced rapid changes in translation efficiency of many genes compared to mock, with more genes that are translationally induced in an injectisome-dependent manner (Figure S4B). Furthermore, comparison of WT *Salmonella* and injectisome mutant infections revealed that injectisome-specific translational upregulation precedes the transcriptional regulation, with relatively little change in mRNA abundances within the first 120 min (Figure 3C-E). Many components of the classical inflammasome that are activated by *Salmonella*^70^, including *Nlrp3*, *Casp1* and *Gsdmd*, are among those transcriptionally upregulated after 120 min of infection with either strain of *Salmonella* (Figure S4C). This is after the initial wave of cell death (Figure 1C) that is typically attributed to inflammasome activation.

Gene ontology enrichment analysis of transcripts with differential translational efficiencies during the first 60 min of WT vs injectisome mutant *Salmonella* infection showed they have functions related to cytokine activity and, strikingly, DNA binding and transcription (Figure 3F, S4D, Table S3). In contrast, and similar to previous transcriptomic studies^3^, the transcriptional response was also enriched for cytokines and other cell signalling genes but there was no such enrichment for DNA binding/transcription related genes. Beyond the initial 60 min, genes with injectisome-dependent transcriptional induction were enriched for DNA binding functions, though the overlap with the rapidly transcriptionally regulated DNA binding factors was small (Figure S4E). Overall, this reinforces the hypothesis that rapid, injectisome-dependent translational induction of transcription modulators reshapes the transcriptional landscape, and consequently, response of macrophages to *Salmonella* infection.

## Discussion

Cellular stress alters gene expression dynamics^1–5,71,72^. Here we show that bacterial infection is a potent stressor that induces selective host protein synthesis through modulating mRNA translation efficiency within the first hour of infection. Much of this regulation of translation efficiency is rapidly triggered during the interaction between the *Salmonella* injectisome and the macrophage plasmalemma, leading to the rapid synthesis of transcriptional modulators. This appears to be triggered predominately by injectisome penetration, rather than the subsequent injection of effector molecules. This study highlights the importance of cellular responses to pathogen attack, and potentially other insults, to be able to rapidly generate proteins, especially DNA binding proteins such as transcription factors, by modulating the translation efficiency of mRNA molecules in the cytoplasm. This provides a mechanism to synergize the transcriptional response to biotic stress.

We found that, in macrophages, one of the mRNAs controlled at the translational level encoded the transcription factor EGR1. EGR1 induction was shown to negatively regulate inflammation and cell death-associated genes resulting in enhanced cell survival and a limited inflammatory response. EGR1 was recently found to be involved in macrophage development by limiting the accessibility of inflammatory gene enhancers through recruitment of the NuRD chromatin remodelling machinery^57^ and is known to be rapidly and transiently induced in response to various stimuli and has been assigned a diverse range of roles, including regulating replication and cell death^54,58,59,73^. Here, we revealed that accumulation of EGR1 protein is increased by a co-ordinated upregulation of translation as well as transcription. Therefore, overall synthesis of EGR1 protein is particularly rapid and robust and occurs within minutes after the bacterial injectisome interacts with the macrophage plasmalemma. The *Egr1* mRNA and EGR1 protein levels both show tight temporal control, with maximal levels at 60 and 120 min, respectively, declining to undetectable levels within 4 h post-infection. This transient induction of EGR1 protein actively contributes to macrophage survival, as supported by our infection study with the EGR1^KO^ mutant. We subsequently revealed that EGR1 acts as a transcriptional suppressor for genes associated with immune processes, including inflammatory genes such as IL-1β, demonstrating the critical role of EGR1 in restraining the immune response during *Salmonella* infection. Their increased expression likely contributes to the observed increases in the rate of death of EGR1^KO^ mutant macrophages following infection. The rapid death of *Salmonella*-infected macrophages is typically attributed to SPI-1^24,25^. Our data suggest that EGR1 restrains pro-inflammatory signals in WT macrophages during *Salmonella* infection, and thereby inhibits cell death^74,75^. While restraining inflammation likely contributes to survival of macrophages, the decrease of pro-inflammatory and pro-cell death gene expression due to transient EGR1 production appears to ultimately but inadvertently benefit the bacterial invader as evidenced by the fact most macrophages that survive injectisome-mediated infection continue to harbour viable bacteria intracellularly.

Our data obtained with *Salmonella* cells expressing the SptP-GFP blocking substrate to inhibit T3SS-mediated effector secretion supports a model where *Egr1* upregulation at the translational and transcriptional levels is triggered by penetration of the host cell membrane by the bacterial injectisome. Recently, a model has been proposed that describes transcriptional induction following exposure to the *Candida albicans* pore forming toxin candidalysin^76^. We inferred that the injectisome-dependent translation and transcription of *Egr1* is activated through a similar mechanism, as supported by observations that other bacterial secretion systems require EGFR and ERK kinases for *Egr1* expression^60,62^.

Supporting this, we show that induction of SPI-1-dependent *Egr1* expression occurs rapidly as *Salmonella* establishes its infection, in the same timeframe as the T3SS is penetrating the macrophages. We show that it is the penetration by the injectisome of the macrophage plasmalemma that is responsible for increased EGR1 expression, not the secretion of effectors.

In summary, we have demonstrated the significance of rapid, transient reprogramming of gene expression, which is mediated primarily by increases in translation of mRNAs enriched for DNA binding proteins. Many of these are transcription factors and will therefore subsequently reshape the transcriptional landscape during the initial hour of *Salmonella* infection of macrophages. Upon encountering host macrophages, *Salmonella* swiftly boosts the expression of SPI-1 structural components, preparing for infection. Penetration of the macrophage membrane by the SPI-1 injectisome is a major trigger for the changes in host gene expression, which leads to rapid and robust protein production including that of EGR1. EGR1 is a transcriptional suppressor of immune genes and therefore transient expression of EGR1 restrains the inflammatory response and host cell death. We hypothesize that *Salmonella* exploits this brief period of immunosuppression to establish infection, leading to later downregulation of cytokine expression and host survival (Figure 4). In conclusion, this work underscores the critical role of translational regulation in defining the response to bacterial pathogens and the importance of Type III injectisome penetration of host cell membranes, a neglected but crucial aspect of the host-bacterial interaction.

**Fig 4:**
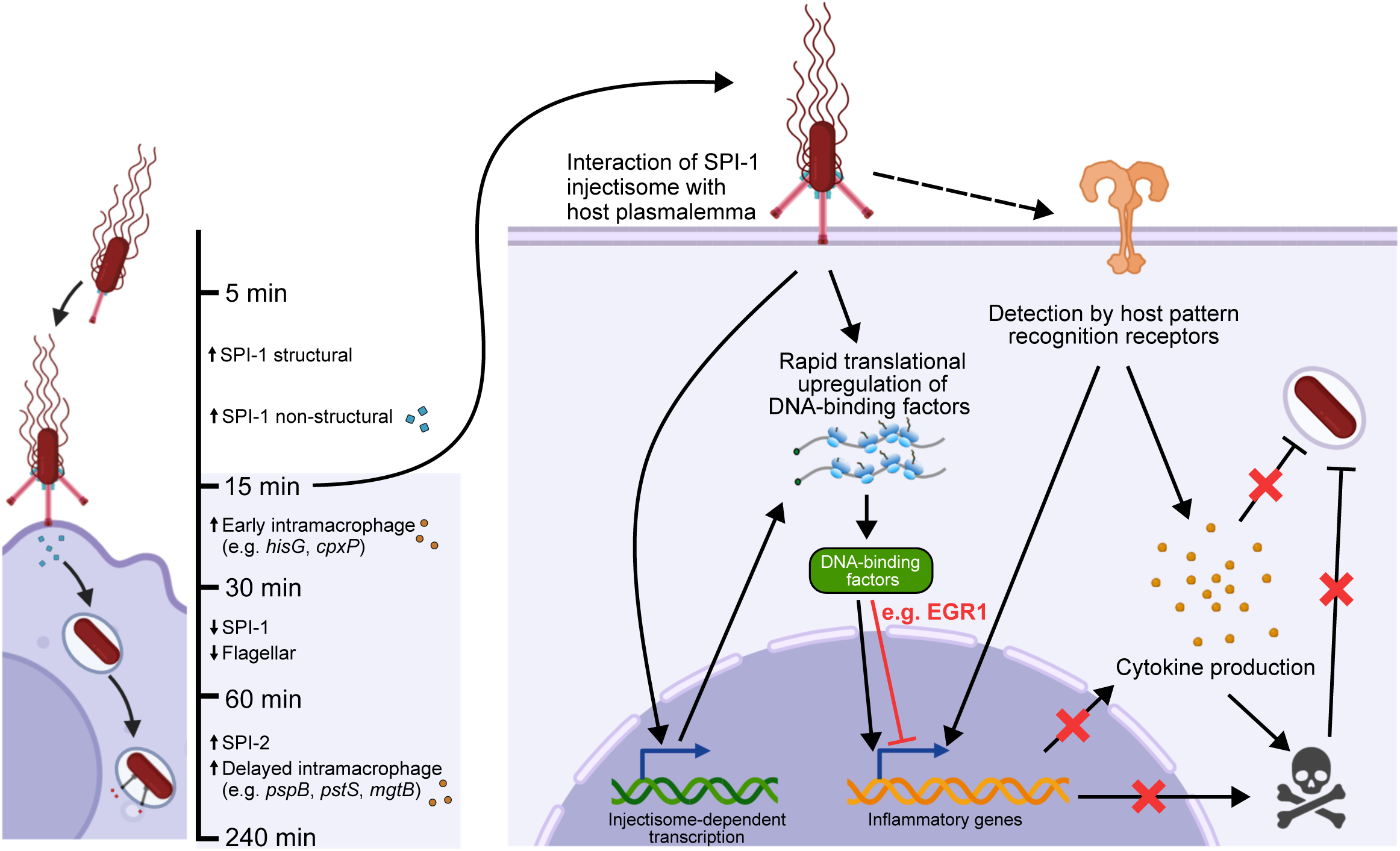
Summary and model. Specific classes of *Salmonella* genes are transiently expressed as infection is established, including rapid upregulation of SPI-1 genes as bacteria encounter host macrophages, leading to increased production of T3SS injectisomes as shown. *Salmonella* SPI-1 injectisome assembly forms transient pores in the host membrane which activates host signaling pathways, leading to increased *Egr1* transcription and enhanced translation. EGR1 negatively modulates immune gene transcription and, in doing so, inflammatory cytokine production in response to *Salmonella* infection is restrained, as is host cell death.

## Methods

### Macrophage cell culture

Primary bone marrow derived macrophages (BMDM) were harvested as previously described^33^. Briefly, bone marrow from the rear legs of C57BL/6 mice was extracted, suspended in culture media, and plated at a density of 10^6^ cells/ml supplemented with 20 ng/ml M-CSF (Peprotech). Cells were differentiated for 7 days, with additional M-CSF supplementation 4 days post-extraction. Immortalized bone marrow-derived macrophages (iBMDM) were generated by retroviral transformation of bone marrow-derived macrophages as previously described^77^. Cells were routinely grown in DMEM supplemented with 10% foetal bovine serum at 37°C, 5% CO_2_.

### Salmonella infection

*Salmonella* Typhimurium SL1344 WT and Δ*prgJ* were sub-cultured in LB from stationary phase cultures and grown at 37°C, with shaking at 200 rpm, until late exponential phase. Bacteria were washed and diluted in culture media and added to cells at a multiplicity of infection of 10. For infections using primary bone marrow-derived macrophages, all media was supplemented with 20 ng/ml M-CSF. Unless otherwise indicated, after 15 min media was supplemented with 100 μg/ml gentamicin.

### Microscopy

At the indicated timepoint, cells were fixed in 4% paraformaldehyde, permeabilized and stained with Phalloidin-CF594 conjugate (Biotium), and goat anti-*Salmonella* CSA-1 (Insight Biotechnology) followed by anti-goat IgG conjugated to Alexa488 (Abcam). Cells were imaged and the proportion infected determined. Macrophages with intracellular bacteria, as determined by the intensity of CSA-1 staining and the position of *Salmonella* within the cell, were considered infected. Macrophages that were uninfected (no intracellular bacteria) but had *Salmonella* associated with the host cell surface membrane were considered to be in the process of becoming infected.

### Gentamicin protection assay

The use of gentamicin protection assay to assess the number of intracellular bacterial has been described previously^2^. This protocol was modified to better account for host cell death. Briefly, 1 h post gentamicin treatment (i.e. 75 min post infection) macrophages were trypsinized, counted, and lysed in 0.09% Triton X-100. Serial dilutions of lysates were plated on LB-agar and grown at 37°C overnight. The *Salmonella* colonies were counted and divided by the number of counted host macrophages.

### Cytotoxicity assay

Cytotoxicity was measured by lactate dehydrogenase release using the CytoTox 96 Non-Radioactive Cytotoxicity Assay kit (Promega) according to the manufacturer’s instructions. Cytotoxicity was determined relative to total cell lysis by 0.9% Triton X-100.

### Ribosome profiling with parallel RNA sequencing

Ribosome profiling was performed as previously described^36^, a schematic of which is presented in Figure 1E. Briefly, at the indicated timepoint, culture supernatant was removed from cells and flash frozen. Cells were lysed in buffer containing cycloheximide and chloramphenicol, and lysates were split for RNA-Seq and Ribo-Seq. For Ribo-Seq, lysates were treated with RNase I and fragments protected from digestion by the ribosome were purified. For RNA-Seq, total cellular RNA was fragmented by alkaline hydrolysis. This was followed by library generation as previously described^36,78–80^. Sequencing was performed using NextSeq-500 or 2000 (Illumina).

Reads were aligned sequentially to mouse rRNA, mouse mRNA, *Salmonella* rRNA and *Salmonella* mRNA. Mouse reference sequences were based on NCBI release mm10, and *Salmonella* reference sequences were based on GenBank sequences FQ312003.1, HE654725.1, HE654726.1 and HE654724.1. RiboSeqR^36^ was used to confirm the quality of libraries and to count reads aligning to coding sequences. Xtail^81^ was used to determine differential translational efficiency of these coding sequences and edgeR^82^ was used for normalization and to filter *Salmonella* genes by expression for retention in further analysis. Gene set enrichment analysis was performed using g:Profiler^83^.

### Western immunoblotting

Protein was harvested from cells disrupted with lysis buffer (50 mM Tris-HCl pH 8.0, 150 mM NaCl, 1 mM EDTA, 10% glycerol, 1% Triton X-100, 0.1% IGEPAL-CA630) containing protease (cOmplete Mini, Roche) and phosphatase inhibitors (PhosSTOP, Roche). 30 µg protein was separated by SDS-PAGE and transferred to nitrocellulose membrane. EGR1 was detected using rabbit anti-EGR1 (Cell Signaling) followed by anti-rabbit conjugated to HRP (Cell Signaling). The HRP signal was assayed using SuperSignal West Pico PLUS substrate (Thermo Scientific). GAPDH was detected using mouse anti-GAPDH (Sigma Aldrich) followed by anti-mouse conjugated to IRDye 800 CW (Licor). SipC was detected by mouse anti-SipC (tgcBIOMICS) followed by anti-mouse IgG antibody conjugated to HRP (Promega). GroEL was detected by rabbit anti-GroEL (Abcam) followed by anti-rabbit IgG antibody conjugated to HRP (Promega). Myc tagged SptP-GFP blocking substrates were detected with anti-Myc-HRP conjugate mouse antibody (Cell Signaling).

### Reverse transcription quantitative polymerase chain reaction (RT-qPCR)

RNA was extracted from cells using TRIzol (Invitrogen) per the manufacturer’s instructions. Reverse transcription was performed using M-MLV reverse transcriptase (Promega) with random hexamer primers ’Promega) per the manufacturer’s instructions. Realtime qPCR was performed using iTaq Universal SYBR Green Supermix (Bio-Rad) and assayed on ViiA 7 system (Applied Biosystems) per the manufacturer’s instructions. Primers were designed using PrimerBLAST (Table S4).

### Knockout of EGR1

The Alt-R CRISPR-Cas system (IDT) was used per the manufactures instructions to edit the *Egr1* coding sequence using guide RNAs targeting *Egr1* or no genes as a negative control (Table S5)^84^. The system was delivered by lipofection into macrophages using Lipofectamine CRISPRMAX (Invitrogen). A clonal population was generated and targeted Sanger sequencing at the *Egr1* locus was performed (Genewiz) to confirm mutation of *Egr1*.

### Blocked injectisome *Salmonella* transformant

Type III injectisomes can be blocked by fusing GFP to the C-terminus of an effector protein^63^. To generate our inducible blocking substrate construct, gDNA encoding the *Salmonella* chaperone SicP (residues 1-116) up to and including the downstream gene encoding the effector protein SptP (residues 1-543) was inserted into the pTrc99a plasmid^85^ in-frame with sequence encoding C-terminal GFP followed by a myc-tag. IPTG induction results in the production of an mRNA transcript encoding wild type SicP chaperone which promotes efficient targeting of SicP’s cognate substrate (in this case the SptP-GFP-myc blocking substrate) to the injectisome export machinery. The mRNA transcript also encodes the SptP-GFP-myc fusion protein which is targeted to the injectisome export machinery and stalls within the export channel, blocking the secretion of effector proteins via the SPI-1 injectisome. To block effector protein secretion via the injectisome, *Salmonella* cells carrying the inducible blocking substrate construct were grow in LB containing 100 μg.ml ampicillin and expression of the blocking substrate (and the SicP chaperone) was achieved by supplementing the media with isopropyl β-D-1-thiogalactopyranoside (IPTG) to a final concentration of 100 μM.

### mRNA 3’ end sequencing

Preparation of mRNA 3’ end sequencing libraries was performed using QuantSeq 3’ mRNA-Seq Library Prep Kit (Lexogen) with TRIzol extracted RNA and libraries were sequenced by Novogene Ltd. Reads were aligned to the mouse genome (mm10) and those aligning to genes counted. Read count normalization and differential expression analysis was performed using edgeR.

### Enzyme-linked immunosorbent assay (ELISA)

Culture supernatants were removed from infected cells at the indicated timepoint. ELISAs were performed to quantify IL-1β in these culture supernatants using the Mouse IL-1 beta/IL-1F2 DuoSet ELISA kit (R&D Systems) per the manufacturer’s instructions.

### Protein export assays

Export assays were performed as previously described^86^. Briefly, *Salmonella* strains were cultured at 37 °C in LB broth with 100 μM IPTG to mid-log phase (OD600nm 1.5) for 2 h. Cells were centrifuged (6000 x g, 3 min) and resuspended in fresh media and grown for a further 60 min at 37 °C. The cells were pelleted by centrifugation (16,000 x g, 5 min) and the supernatant passed through a 0.2 μm nitrocellulose filter. Proteins were precipitated with 10% trichloroacetic acid (TCA) and 1% Triton X-100 on ice for 1 hr, pelleted by centrifugation (16,000 x g, 10 min), washed with ice-cold acetone, and resuspended in SDS-PAGE loading buffer (volumes calibrated according to cell densities). Fractions were analyzed by immunoblotting with anti-SipC (tgcBIOMICS), anti-Myc (Cell Signaling) and anti-GroEL (Abcam) anti-sera.

## Data availability

Raw and processed data are available from ArrayExpress accessions E-MTAB-13212 and E-MTAB-13213 or can be found in the supplementary tables. Customized scripts used for this project are available upon request.

## Acknowledgements

We would also like to thank Jim Kaufman, Klaus Okkenhaug, and Alex Murphy for discussions and comments on the manuscript. F.L. was supported by a BBSRC DTP studentship. G.W. was supported by the Department of Pathology PhD studentship, B.Y.W.C., R.J. and M.B. and O.J.B. are supported by a Medical Research Council Fellowship to B.Y.W.C. [MR/R021821/1]. J.P. and M.B. are supported by a BBSRC project grant awarded to B.Y.W.C. [BB/V006096/1]. The B.Y.W.C. laboratory is supported by a Medical Research Council Fellowship [MR/R021821/1], BBSRC project grants [BB/X001261/1, BB/V017780/1 and BB/V006096/1] and a Royal Society Research Grant [RGS\R2\192222]. Figures created with BioRender.com.

## Author contributions

B.Y.W.C. conceived the research. B.Y.W.C., G.W., R.J., J.P., O.J.B., F.L., J.P.C. and C.B. designed experiments. R.J., J.P. and B.Y.W.C. generated RiboSeq, RNA-Seq and QuantSeq libraries. G.W., R.J. and J.P. performed molecular and cell biology experiments. O.J.B. generated *Salmonella* mutants. F.L. performed bacterial bioinformatics. B.Y.W.C., G.W., R.J., J.P., F.L. and M.B. performed the bioinformatic analysis. C.B. provided macrophages and *prgJ* mutant. P.T. provided training for extracting BMDM. B.Y.W.C., G.W., O.J.B. and J.P.C. wrote the manuscript.

---

**Fig S1:** (**A**) Percentage of macrophages infected by WT *Salmonella* determined by microscopy at 15 and 60 min, with and without addition of gentamicin at 15 min. Significance determined by Student’s t-test; *n*=2. (**B**) Percentage of macrophages infected by WT or Δ*prgJ Salmonella* determined by microscopy at 15 min. Significance determined by Student’s t-test; *n*=2. (**C**) Meta-gene translatome from ribosome profiling of *Salmonella* infected macrophages at 60 min. Histograms of RPF 5′ ends relative to start and stop codons colored by their reading frame relative to the coding sequence. (**D**) Heatmap showing mRNA abundance, protein synthesis and TE of all genes in Fig 1G. Genes are ordered by hierarchical clustering of mRNA abundance across all conditions (left). Arrows indicate direction of differential transcript abundance (log_2_FC ±1) in the indicated infection vs mock.(**E**) Ribo-Seq and RNA-Seq transcript coverage of *Atf4*. Ribo-Seq reads are represented by their P site position and colored by their reading frame relative to the start codon of the main *Atf3* coding sequence. Open reading frames are represented by bars above each plot. (**F**) Comparison of changes in mRNA abundance (left) and TE (right) on infection with WT over mock infection or Δ*prgJ Salmonella*. Genes upregulated transcriptionally in both (log_2_FC > 1) are shown in red. *Egr1*, *Nr4a1* and *Nfkbiz* are plotted separately due to low abundance in mock infection precluding accurate calculation of TE, and as such TE fold change in WT *Salmonella* over mock infection. (**F**) Normalised mRNA abundance and protein synthesis of *Nlrp3* at 60 min post-infection. (**G**) Ribo-Seq and RNA-Seq transcript coverage of *Nlrp3* as in **E**. The *Nlrp3* uORF can be readily seen as reads in a different reading frame within the 5’ UTR.

**Fig S2:** (**A**) Example section of the *Egr1* coding sequence with an alignment from targeted sequencing of the Egr1 locus in the EGR1 KO macrophages. Mismatched bases and deletions are highlighted in red. (**B**) Principal component analysis of the transcriptomes of WT *Salmonella* infected or mock infected EGR1 KO or control macrophages over an infection time course.

**Fig S3:** (**A**) Percentage of primary macrophages infected by WT or Δ*prgJ Salmonella* determined by microscopy at 15 min. (**B**) Meta-gene translatome from ribosome profiling across a *Salmonella* primary bone marrow derived macrophage infection time course. Histograms of RPF 5′ ends relative to start and stop codons colored by their reading frame relative to the coding sequence.

**Fig S4:** (**A**) log_2_FC TE vs log_2_FC mRNA abundance between the indicated infections, across the time course. Dashed orange lines show the log_2_FC cutoffs (±2) used to select differentially expressed genes; genes that pass these thresholds are colored. (**B**) Number of genes differentially expressed between WT vs mock and Δ*prgJ* vs mock infection at both the TE (right) and mRNA (left) levels. (**C**) Expression of genes encoding components of the inflammasome^70^ that are upregulated over the infection timecourse. (**D**) Top 10 enriched GO molecular function terms in differentially expressed genes at both the TE (right) and mRNA (left) levels. Genes were split by when they were differentially expressed: at or before 60 min, and after 60 min post infection. (**E**) Overlap of genes with DNA binding and transcription related annotations that are differentially regulated on the translational level at or before 60 min or the transcriptional level after 60 min in the comparison of WT vs Δ*prgJ* infection.

## References

1. Denzer, L., Schroten, H., and Schwerk, C. (2020). From gene to protein - how bacterial virulence factors manipulate host gene expression during infection. Int. J. Mol. Sci. 21, 3730. 10.3390/ijms21103730.

2. Hannemann, S., Gao, B., and Galán, J.E. (2013). Salmonella modulation of host cell gene expression promotes its intracellular growth. PLoS Pathog. 9, e1003668. 10.1371/journal.ppat.1003668.

3. Jensen, K., Gallagher, I.J., Kaliszewska, A., Zhang, C., Abejide, O., Gallagher, M.P., Werling, D., and Glass, E.J. (2016). Live and inactivated Salmonella enterica serovar Typhimurium stimulate similar but distinct transcriptome profiles in bovine macrophages and dendritic cells. Vet. Res. 47. 10.1186/s13567-016-0328-y.

4. Ordas, A., Hegedus, Z., Henkel, C. V, Stockhammer, O.W., Butler, D., Jansen, H.J., Racz, P., Mink, M., Spaink, H.P., and Meijer, A.H. (2011). Deep sequencing of the innate immune transcriptomic response of zebrafish embryos to Salmonella infection. Fish Shellfish Immunol. 31, 716–724. 10.1016/j.fsi.2010.08.022.

5. Maekawa, S., Wang, P.C., and Chen, S.C. (2019). Comparative study of immune reaction against bacterial infection from transcriptome analysis. Front. Immunol. 10. 10.3389/fimmu.2019.00153.

6. Westermann, A.J., Venturini, E., Sellin, M.E., Förstner, K.U., Hardt, W.D., and Vogel, J. (2019). The major RNA-binding protein ProQ impacts virulence gene expression in Salmonella enterica serovar Typhimurium. MBio 10, e02504–2518. 10.1128/mBio.02504-18.

7. Fields, P.I., Swanson, R. V., Haidaris, C.G., and Heffron, F. (1986). Mutants of Salmonella Typhimurium that cannot survive within the macrophage are avirulent. PNAS 83, 5189–5193. 10.1073/pnas.83.14.5189.

8. Liss, V., Swart, A.L., Kehl, A., Hermanns, N., Zhang, Y., Chikkaballi, D., Böhles, N., Deiwick, J., and Hensel, M. (2017). Salmonella enterica remodels the host cell endosomal system for efficient intravacuolar nutrition. Cell Host Microbe 21, 390–402. 10.1016/j.chom.2017.02.005.

9. Jiang, L., Wang, P., Song, X., Zhang, H., Ma, S., Wang, J., Li, W., Lv, R., Liu, X., Ma, S., et al. (2021). Salmonella Typhimurium reprograms macrophage metabolism via T3SS effector SopE2 to promote intracellular replication and virulence. Nat. Commun. 12, 1–18. 10.1038/s41467-021-21186-4.

10. Miletic, S., Fahrenkamp, D., Goessweiner-Mohr, N., Wald, J., Pantel, M., Vesper, O., Kotov, V., and Marlovits, T.C. (2021). Substrate-engaged type III secretion system structures reveal gating mechanism for unfolded protein translocation. Nat. Commun. 12, 1546. 10.1038/s41467-021-21143-1.

11. Myeni, S.K., Wang, L., and Zhou, D. (2013). SipB-SipC complex is essential for translocon formation. PLoS One 8, e60499. 10.1371/journal.pone.0060499.

12. Miki, T., Okada, N., Shimada, Y., and Danbara, H. (2004). Characterization of Salmonella pathogenicity island 1 type III secretion-dependent hemolytic activity in Salmonella enterica serovar Typhimurium. Microb. Pathog. 37, 65–72. 10.1016/j.micpath.2004.04.006.

13. Ly, K.T., and Casanova, J.E. (2007). Mechanisms of Salmonella entry into host cells. Cell. Microbiol. 9, 2103–2111. 10.1111/j.1462-5822.2007.00992.x.

14. Galán, J.E. (1998). Interactions of Salmonella with host cells: Encounters of the closest kind. Proc. Natl. Acad. Sci. 95, 14006–14008. 10.1073/pnas.95.24.14006.

15. Hume, P.J., Singh, V., Davidson, A.C., and Koronakis, V. (2017). Swiss army pathogen: The Salmonella entry toolkit. Front. Cell. Infect. Microbiol. 7, 348. 10.3389/fcimb.2017.00348.

16. Vazquez-Torres, A., Xu, Y., Jones-Carson, J., Holden, D.W., Lucia, S.M., Dinauer, M.C., Mastroeni, P., and Fang, F.C. (2000). Salmonella Pathogenicity Island 2-Dependent Evasion of the Phagocyte NADPH Oxidase. Science (80-.). 287, 1655–1658. 10.1126/science.287.5458.1655.

17. Guignot, J., and Tran Van Nhieu, G. (2016). Bacterial control of pores induced by the type III secretion system: mind the gap. Front. Immunol. 7, 84. 10.3389/fimmu.2016.00084.

18. Park, D., Lara-Tejero, M., Waxham, M.N., Li, W., Hu, B., Galán, J.E., and Liu, J. (2018). Visualization of the type III secretion mediated salmonella–host cell interface using cryo-electron tomography. Elife 7. 10.7554/ELIFE.39514.

19. Murthy, S., Karkossa, I., Schmidt, C., Hoffmann, A., Hagemann, T., Rothe, K., Seifert, O., Anderegg, U., von Bergen, M., Schubert, K., et al. (2022). Danger signal extracellular calcium initiates differentiation of monocytes into SPP1/osteopontin-producing macrophages. Cell Death Dis. 2022 131 13, 1–15. 10.1038/s41419-022-04507-3.

20. Koumangoye, R. (2022). The role of Cl-and K+efflux in NLRP3 inflammasome and innate immune response activation. Am. J. Physiol. -Cell Physiol. 322, C645–C652. 10.1152/AJPCELL.00421.2021/ASSET/IMAGES/LARGE/AJPCELL.00421.2021_F002.JPEG.

21. Richter-Dahlfors, A., Buchan, A.M.J., and Finlay, B.B. (1997). Murine salmonellosis studied by confocal microscopy: Salmonella Typhimurium resides intracellularly inside macrophages and exerts a cytotoxic effect on phagocytes in vivo. J. Exp. Med. 186, 569–580. 10.1084/jem.186.4.569.

22. Price, J. V., and Vance, R.E. (2014). The macrophage paradox. Immunity 41, 685– 693. 10.1016/j.immuni.2014.10.015.

23. Talbot, S., Tötemeyer, S., Yamamoto, M., Akira, S., Hughes, K., Gray, D., Barr, T., Mastroeni, P., Maskell, D.J., and Bryant, C.E. (2009). Toll-like receptor 4 signalling through MyD88 is essential to control Salmonella enterica serovar Typhimurium infection, but not for the initiation of bacterial clearance. Immunology 128, 472. 10.1111/J.1365-2567.2009.03146.X.

24. Grant, A.J., Sheppard, M., Deardon, R., Brown, S.P., Foster, G., Bryant, C.E., Maskell, D.J., and Mastroeni, P. (2008). Caspase-3-dependent phagocyte death during systemic Salmonella enterica serovar Typhimurium infection of mice. Immunology 125, 28–37. 10.1111/j.1365-2567.2008.02814.x.

25. Gram, A.M., Wright, J.A., Pickering, R.J., Lam, N.L., Booty, L.M., Webster, S.J., and Bryant, C.E. (2021). Salmonella flagellin activates NAIP/NLRC4 and canonical NLRP3 inflammasomes in human macrophages. J. Immunol. 206, 631–640. 10.4049/jimmunol.2000382.

26. Bergsbaken, T., Fink, S.L., and Cookson, B.T. (2009). Pyroptosis: host cell death and inflammation. Nat. Rev. Microbiol. 7, 99–109. 10.1038/nrmicro2070.

27. Chen, L.M., Kaniga, K., and Galán, J.E. (1996). Salmonella spp. are cytotoxic for cultured macrophages. Mol. Microbiol. 21, 1101–1115. 10.1046/j.1365-2958.1996.471410.x.

28. Monack, D.M., Raupach, B., Hromockyj, A.E., and Falkow, S. (1996). Salmonella Typhimurium invasion induces apoptosis in infected macrophages. PNAS 93, 9833– 9838. 10.1073/pnas.93.18.9833.

29. Lundberg, U., Vinatzer, U., Berdnik, D., von Gabain, A., and Baccarini, M. (1999). Growth phase-regulated induction of Salmonella-induced macrophage apoptosis correlates with transient expression of SPI-1 genes. J. Bacteriol. 181, 3433–3437. 10.1128/JB.181.11.3433-3437.1999.

30. Wang, L., Trebicka, E., Fu, Y., Waggoner, L., Akira, S., Fitzgerald, K.A., Kagan, J.C., and Cherayil, B.J. (2011). Regulation of lipopolysaccharide-induced translation of tumor necrosis factor-alpha by the toll-like receptor 4 adaptor protein TRAM. J. Innate Immun. 3, 437–446. 10.1159/000324833.

31. Dumitru, C.D., Ceci, J.D., Tsatsanis, C., Kontoyiannis, D., Stamatakis, K., Lin, J.-H., Patriotis, C., Jenkins, N.A., Copeland, N.G., Kollias, G., et al. (2000). TNF-α Induction by LPS Is Regulated Posttranscriptionally via a Tpl2/ERK-Dependent Pathway. Cell 103, 1071–1083. 10.1016/S0092-8674(00)00210-5.

32. Drecktrah, D., Knodler, L.A., Ireland, R., and Steele-Mortimer, O. (2006). The mechanism of Salmonella entry determines the vacuolar environment and intracellular gene expression. Traffic 7, 39–51. 10.1111/J.1600-0854.2005.00360.X.

33. Man, S.M., Hopkins, L.J., Nugent, E., Cox, S., Glück, I.M., Tourlomousis, P., Wright, J.A., Cicuta, P., Monie, T.P., and Bryant, C.E. (2014). Inflammasome activation causes dual recruitment of NLRC4 and NLRP3 to the same macromolecular complex. Proc. Natl. Acad. Sci. 111, 7403–7408. 10.1073/PNAS.1402911111.

34. Fink, S.L., and Cookson, B.T. (2007). Pyroptosis and host cell death responses during Salmonella infection. Cell. Microbiol. 9, 2562–2570. 10.1111/J.1462-5822.2007.01036.X.

35. Ingolia, N.T., Ghaemmaghami, S., Newman, J.R.S., and Weissman, J.S. (2009). Genome-wide analysis in vivo of translation with nucleotide resolution using ribosome profiling. Science 324, 218–223. 10.1126/SCIENCE.1168978.

36. Chung, B.Y., Hardcastle, T.J., Jones, J.D., Irigoyen, N., Firth, A.E., Baulcombe, D.C., and Brierley, I. (2015). The use of duplex-specific nuclease in ribosome profiling and a user-friendly software package for Ribo-seq data analysis. RNA 21, 1731. 10.1261/RNA.052548.115.

37. Vattem, K.M., and Wek, R.C. (2004). Reinitiation involving upstream ORFs regulates ATF4 mRNA translation in mammalian cells. Proc. Natl. Acad. Sci. 101, 11269– 11274. 10.1073/pnas.0400541101.

38. Landmann, R., Knopf, H.P., Link, S., Sansano, S., Schumann, R., and Zimmerli, W. (1996). Human monocyte CD14 is upregulated by lipopolysaccharide. Infect. Immun. 64, 1762–1769. 10.1128/IAI.64.5.1762-1769.1996.

39. Yang, H., Li, Q., Stroup, E.K., Wang, S., and Ji, Z. (2024). Widespread stable noncanonical peptides identified by integrated analyses of ribosome profiling and ORF features. Nat. Commun. 2024 151 15, 1–18. 10.1038/s41467-024-46240-9.

40. Bryant, C. (2021). Inflammasome activation by Salmonella. Curr. Opin. Microbiol. 64, 27–32. 10.1016/j.mib.2021.09.004.

41. Qu, Y., Misaghi, S., Newton, K., Maltzman, A., Izrael-Tomasevic, A., Arnott, D., and Dixit, V.M. (2016). NLRP3 recruitment by NLRC4 during Salmonella infection. J. Exp. Med. 213, 877–885. 10.1084/JEM.20132234.

42. Li, P., Allen, H., Banerjee, S., Franklin, S., Herzog, L., Johnston, C., McDowell, J., Paskind, M., Rodman, L., Salfeld, J., et al. (1995). Mice deficient in IL-1 beta-converting enzyme are defective in production of mature IL-1 beta and resistant to endotoxic shock. Cell 80, 401–411. 10.1016/0092-8674(95)90490-5.

43. Hersh, D., Monack, D.M., Smith, M.R., Ghori, N., Falkow, S., and Zychlinsky, A. (1999). The Salmonella invasin SipB induces macrophage apoptosis by binding to caspase-1. Proc. Natl. Acad. Sci. 96, 2396–2401. 10.1073/pnas.96.5.2396.

44. Cook, P., Tötemeyer, S., Stevenson, C., Fitzgerald, K.A., Yamamoto, M., Akira, S., Maskell, D.J., and Bryant, C.E. (2007). Salmonella-induced SipB-independent cell death requires Toll-like receptor-4 signalling via the adapter proteins Tram and Trif. Immunology 122, 222. 10.1111/J.1365-2567.2007.02631.X.

45. Reyes Ruiz, V.M., Ramirez, J., Naseer, N., Palacio, N.M., Siddarthan, I.J., Yan, B.M., Boyer, M.A., Pensinger, D.A., Sauer, J.-D., and Shin, S. (2017). Broad detection of bacterial type III secretion system and flagellin proteins by the human NAIP/NLRC4 inflammasome. Proc. Natl. Acad. Sci. 114, 13242–13247. 10.1073/pnas.1710433114.

46. Mariathasan, S., Newton, K., Monack, D.M., Vucic, D., French, D.M., Lee, W.P., Roose-Girma, M., Erickson, S., and Dixit, V.M. (2004). Differential activation of the inflammasome by caspase-1 adaptors ASC and Ipaf. Nature 430, 213–218. 10.1038/nature02664.

47. Martin-Sanchez, F., Diamond, C., Zeitler, M., Gomez, A.I., Baroja-Mazo, A., Bagnall, J., Spiller, D., White, M., Daniels, M.J.D., Mortellaro, A., et al. (2016). Inflammasome-dependent IL-1β release depends upon membrane permeabilisation. Cell Death Differ. 23, 1219–1231. 10.1038/cdd.2015.176.

48. Lau, L.F., and Nathans, D. (1987). Expression of a set of growth-related immediate early genes in BALB/c 3T3 cells: coordinate regulation with c-fos or c-myc. Proc. Natl. Acad. Sci. 84, 1182–1186. 10.1073/PNAS.84.5.1182.

49. McMahon, S.B., and Monroe, J.G. (1996). The role of early growth response gene 1 (egr-1) in regulation of the immune response. J. Leukoc. Biol. 60, 159–166. 10.1002/JLB.60.2.159.

50. Bahrami, S., and Drabløs, F. (2016). Gene regulation in the immediate-early response process. Adv. Biol. Regul. 62, 37–49. 10.1016/J.JBIOR.2016.05.001.

51. Hamers, A.A.J., van Dam, L., Teixeira Duarte, J.M., Vos, M., Marinković, G., van Tiel, C.M., Meijer, S.L., van Stalborch, A.-M., Huveneers, S., te Velde, A.A., et al. (2015). Deficiency of nuclear receptor Nur77 aggravates mouse experimental colitis by increased NFκB activity in macrophages. PLoS One 10, e0133598. 10.1371/journal.pone.0133598.

52. Li, L., Liu, Y., Chen, H., Li, F., Wu, J., Zhang, H., He, J., Xing, Y., Chen, Y., Wang, W., et al. (2015). Impeding the interaction between Nur77 and p38 reduces LPS-induced inflammation. Nat. Chem. Biol. 11, 339–346. 10.1038/nchembio.1788.

53. Partyka, J., Henkel, M., Campfield, B.T., and 14, P.A. (2020). A Novel Role for the Nuclear Receptor, NR4A1, in Klebsiella pneumoniae Lung Infection. bioRxiv, 2020.09.03.282475. 10.1101/2020.09.03.282475.

54. Thiel, G., and Cibelli, G. (2002). Regulation of life and death by the zinc finger transcription factor Egr-1. J. Cell. Physiol. 193, 287–292. 10.1002/jcp.10178.

55. Chbicheb, S., Yao, X., Rodeau, J.-L., Salamone, S., Boisbrun, M., Thiel, G., Spohn, D., Grillier-Vuissoz, I., Chapleur, Y., Flament, S., et al. (2011). EGR1 expression: A calcium and ERK1/2 mediated PPARγ-independent event involved in the antiproliferative effect of 15-deoxy-Δ12,14-prostaglandin J2 and thiazolidinediones in breast cancer cells. Biochem. Pharmacol. 81, 1087–1097. 10.1016/j.bcp.2011.02.006.

56. Banerji, R., and Saroj, S.D. (2021). Early growth response 1 (EGR1) activation in initial stages of host–pathogen interactions. Mol. Biol. Rep. 48, 2935–2943. 10.1007/s11033-021-06305-0.

57. Trizzino, M., Zucco, A., Deliard, S., Wang, F., Barbieri, E., Veglia, F., Gabrilovich, D., and Gardini, A. (2021). EGR1 is a gatekeeper of inflammatory enhancers in human macrophages. Sci. Adv. 7. 10.1126/sciadv.aaz8836.

58. Kimura, T.E., Duggirala, A., Hindmarch, C.C.T., Hewer, R.C., Cui, M.Z., Newby, A.C., and Bond, M. (2014). Inhibition of Egr1 expression underlies the anti-mitogenic effects of cAMP in vascular smooth muscle cells. J. Mol. Cell. Cardiol. 72, 9–19. 10.1016/j.yjmcc.2014.02.001.

59. Sukhatme, V.P., Cao, X., Chang, L.C., Tsai-Morris, C.H., Stamenkovich, D., Ferreira, P.C.P., Cohen, D.R., Edwards, S.A., Shows, T.B., Curran, T., et al. (1988). A zinc finger-encoding gene coregulated with c-fos during growth and differentiation, and after cellular depolarization. Cell 53, 37–43. 10.1016/0092-8674(88)90485-0.

60. de Grado, M., Rosenberger, C.M., Gauthier, A., Vallance, B.A., and Finlay, B.B. (2001). Enteropathogenic Escherichia coli infection induces expression of the early growth response factor by activating mitogen-activated protein kinase cascades in epithelial cells. Infect. Immun. 69, 6217–6224. 10.1128/IAI.69.10.6217-6224.2001.

61. Shin, H., and Cornelis, G.R. (2007). Type III secretion translocation pores of Yersinia enterocolitica trigger maturation and release of pro-inflammatory IL-1β. Cell. Microbiol. 9, 2893–2902. 10.1111/j.1462-5822.2007.01004.x.

62. Keates, S., Keates, A.C., Nath, S., Peek, R.M., and Kelly, C.P. (2005). Transactivation of the epidermal growth factor receptor by cag+ Helicobacter pylori induces upregulation of the early growth response gene Egr-1 in gastric epithelial cells. Gut 54, 1363–1369. 10.1136/gut.2005.066977.

63. Radics, J., Königsmaier, L., and Marlovits, T.C. (2013). Structure of a pathogenic type 3 secretion system in action. Nat. Struct. Mol. Biol. 2013 211 21, 82–87. 10.1038/nsmb.2722.

64. Kudryashev, M., Diepold, A., Amstutz, M., Armitage, J.P., Stahlberg, H., and Cornelis, G.R. (2015). Yersinia enterocolitica type III secretion injectisomes form regularly spaced clusters, which incorporate new machines upon activation. Mol. Microbiol. 95, 875–884. 10.1111/MMI.12908/SUPPINFO.

65. Eriksson, S., Lucchini, S., Thompson, A., Rhen, M., and Hinton, J.C.D. (2003). Unravelling the biology of macrophage infection by gene expression profiling of intracellular Salmonella enterica. Mol. Microbiol. 47, 103–118. 10.1046/j.1365-2958.2003.03313.x.

66. Srikumar, S., Kröger, C., Hébrard, M., Colgan, A., Owen, S. V., Sivasankaran, S.K., Cameron, A.D.S., Hokamp, K., and Hinton, J.C.D. (2015). RNA-seq brings new insights to the intra-macrophage transcriptome of Salmonella Typhimurium. PLOS Pathog. 11, e1005262. 10.1371/journal.ppat.1005262.

67. Chakraborty, S., Mizusaki, H., and Kenney, L.J. (2015). A FRET-based DNA biosensor tracks OmpR-dependent acidification of Salmonella during macrophage infection. PLOS Biol. 13, e1002116. 10.1371/journal.pbio.1002116.

68. Pérez-Morales, D., Banda, M.M., Chau, N.Y.E., Salgado, H., Martínez-Flores, I., Ibarra, J.A., Ilyas, B., Coombes, B.K., and Bustamante, V.H. (2017). The transcriptional regulator SsrB is involved in a molecular switch controlling virulence lifestyles of Salmonella. PLOS Pathog. 13, e1006497. 10.1371/journal.ppat.1006497.

69. Brown, N.F., Rogers, L.D., Sanderson, K.L., Gouw, J.W., Hartland, E.L., and Foster, L.J. (2014). A horizontally acquired transcription factor coordinates Salmonella adaptations to host microenvironments. MBio 5, 1727–1741. 10.1128/mBio.01727-14.

70. Bryant, C. (2021). Inflammasome activation by Salmonella. Curr. Opin. Microbiol. 64, 27–32. 10.1016/j.mib.2021.09.004.

71. Advani, V.M., and Ivanov, P. (2019). Translational control under stress: reshaping the translatome. BioEssays 41, e1900009. 10.1002/BIES.201900009.

72. Hoang, H.D., Neault, S., Pelin, A., and Alain, T. (2021). Emerging translation strategies during virus–host interaction. Wiley Interdiscip. Rev. RNA 12. 10.1002/WRNA.1619.

73. Yan, S.F., Fujita, T., Lu, J., Okada, K., Shan Zou, Y., Mackman, N., Pinsky, D.J., and Stern, D.M. (2000). Egr-1, a master switch coordinating upregulation of divergent gene families underlying ischemic stress. Nat. Med. 6, 1355–1361. 10.1038/82168.

74. Hughes, S.A., Lin, M., Weir, A., Huang, B., Xiong, L., Chua, N.K., Pang, J., Santavanond, J.P., Tixeira, R., Doerflinger, M., et al. (2023). Caspase-8-driven apoptotic and pyroptotic crosstalk causes cell death and I-1β release in X-linked inhibitor of apoptosis (XIAP) deficiency. EMBO J. 42. 10.15252/embj.2021110468.

75. Wang, Y., and Kanneganti, T.-D. (2021). From pyroptosis, apoptosis and necroptosis to PANoptosis: A mechanistic compendium of programmed cell death pathways. Comput. Struct. Biotechnol. J. 19, 4641–4657. 10.1016/j.csbj.2021.07.038.

76. Ponde, N.O., Lortal, L., Tsavou, A., Hepworth, O.W., Wickramasinghe, D.N., Ho, J., Richardson, J.P., Moyes, D.L., Gaffen, S.L., and Naglik, J.R. (2022). Receptor-kinase EGFR-MAPK adaptor proteins mediate the epithelial response to Candida albicans via the cytolytic peptide toxin, candidalysin. J. Biol. Chem. 298. 10.1016/j.jbc.2022.102419.

77. De Nardo, D., Kalvakolanu, D. V., and Latz, E. (2018). Immortalization of Murine Bone Marrow-Derived Macrophages. In Methods in molecular biology (Clifton, N.J.) (Humana Press Inc.), pp. 35–49. 10.1007/978-1-4939-7837-3_4.

78. Irigoyen, N., Firth, A.E., Jones, J.D., Chung, B.Y.W., Siddell, S.G., and Brierley, I. (2016). High-resolution analysis of Coronavirus gene expression by RNA sequencing and ribosome profiling. PLOS Pathog. 12, e1005473. 10.1371/journal.ppat.1005473.

79. Chung, B.Y.W., Deery, M.J., Groen, A.J., Howard, J., and Baulcombe, D.C. (2017). Endogenous miRNA in the green alga Chlamydomonas regulates gene expression through CDS-targeting. Nat. Plants 2017 310 3, 787–794. 10.1038/s41477-017-0024-6.

80. Chung, B.Y.W., Balcerowicz, M., Di Antonio, M., Jaeger, K.E., Geng, F., Franaszek, K., Marriott, P., Brierley, I., Firth, A.E., and Wigge, P.A. (2020). An RNA thermoswitch regulates daytime growth in Arabidopsis. Nat. Plants 2020 65 6, 522–532. 10.1038/s41477-020-0633-3.

81. Xiao, Z., Zou, Q., Liu, Y., and Yang, X. (2016). Genome-wide assessment of differential translations with ribosome profiling data. Nat. Commun. 7. 10.1038/ncomms11194.

82. Robinson, M.D., McCarthy, D.J., and Smyth, G.K. (2010). edgeR: a Bioconductor package for differential expression analysis of digital gene expression data. Bioinformatics 26, 139–140. 10.1093/BIOINFORMATICS/BTP616.

83. Peterson, H., Kolberg, L., Raudvere, U., Kuzmin, I., and Vilo, J. (2020). gprofiler2 - an R package for gene list functional enrichment analysis and namespace conversion toolset g:Profiler. F1000Research 9, 709. 10.12688/f1000research.24956.2.

84. Wise, D. (2019). Understanding antigen processing in chickens using genome editing technology. 10.17863/CAM.40666.

85. Amann, E., Ochs, B., and Abel, K.J. (1988). Tightly regulated tac promoter vectors useful for the expression of unfused and fused proteins in Escherichia coli. Gene 69, 301–315. 10.1016/0378-1119(88)90440-4.

86. Bryant, O.J., Dhillon, P., Hughes, C., and Fraser, G.M. (2022). Recognition of discrete export signals in early flagellar subunits during bacterial type III secretion. Elife 11. 10.7554/eLife.66264.

